# Viability Differentiation Improves the Diagnostic Potential of 16S rRNA Gene Sequencing in Ready-to-Eat Meat Manufacturing

**DOI:** 10.1101/2025.04.16.649186

**Authors:** Jessica A. Brown, Steven C. Ricke

## Abstract

Molecular-based microbiological approaches have become valuable tools for the food industry. However, even the most advanced molecular techniques are limited in their ability to differentiate based on viability creating the potential for biased results when applied to the food industry. The objective of this study was to generate a viable microbial bio-map of a commercial ready-to-eat (RTE) meat manufacturing process and assess its utility as a diagnostic tool. Product samples were collected from a commercial RTE meat manufacturing facility at various locations throughout processing. Samples were homogenized and aliquoted for culture-based microbial isolation and 16S rRNA gene sequencing. Homogenates were split into pairs and subject to either no treatment (Control) or treated with 25 μM PMAxx (PMA) to remove free and non-viable cellular DNA. Overall, PMA treatment resulted in a less rich microbial community compared to Control samples. Paired analysis revealed that the impact of PMA varied by location with the greatest effects being observed at the beginning and end of manufacturing. Both Control and PMA treated samples identified a shift in the microbial population after thermal processing; however, only PMA treated samples identified a secondary shift in the microbial population occurring after slicing. Taxonomic analysis identified *Lactobacillus* as a predominant genera in sliced and packaged products. These results were further confirmed by the identification of *Lactobacillus sakei* on packaged product using a culture-based approach. These results suggest PMA treatment provides a higher level of sequencing resolution by removing background DNA.

**Importance:** Microbial bio-mapping is a valuable tool for the meat and poultry industry to assess process control and evaluate the efficacy of intervention systems. In recent years it has become more common to incorporate the use of molecular techniques, such as qPCR and 16S rRNA, to quantitatively track target pathogens and gain a more holistic understanding of the microbial community throughout processing. One major limitation we face when applying these DNA-based techniques to the food industry is their inability to differentiate between DNA from viable versus non-viable cells, which may result in the false identification of pathogenic or spoilage microorganisms and bias microbiota results. To practically apply this technology in a ready-to-eat meat manufacturing setting, it is crucial to develop and validate strategies that are capable of differentiating between viable and non-viable cellular DNA.

## Introduction

Food safety and quality continue to be an area of high priority for the meat and poultry industry (Grunert, 2005). The U.S. Department of Agriculture Food Safety and Inspection Service (USDA FSIS) is responsible for ensuring that meat, poultry, and egg products are safe, wholesome and accurately labeled. While deviations in product quality pose little risk to consumer health, it is a major cause of food loss and waste worldwide. At the retail and consumer level alone, it is estimated that the economic cost of wasted meat, poultry, and fish products is $48.5 billion per year (Buzby et al., 2014). Product quality is the culmination of a variety of factors including, but not limited to physical characteristics, production practices, and nutritional content. Together these factors impact overall consumer perception and, in turn, economic value of all food product (Petrescu et al., 2019). From a microbiological perspective, spoilage poses a major threat to lasting product quality. The voluntary process of date marking can help consumers and retailers decide when food is at its highest quality; however, supply chain deviations such as contamination during processing, packaging malfunctions, and improper temperature control, can all result in the premature spoilage of food products and potential damage to a brands reputation (9 CFR 317.8) (van Herpen & de Hooge, 2019). Spoilage is the result of complex interactions between microorganisms and the environment, often resulting in biochemical changes that negatively impact product color, texture, taste, and odor (Erkmen & Bozoglu, 2016). The relatively small group of microorganisms responsible for causing spoilage in meat and poultry products are well-documented and include genera such as lactic acid bacteria, *Pseudomonas*, and *Brochothrix*, all producing a variety of spoilage phenotypes such as slime, discoloration, and gas production (Borch et al., 1996; Erkmen & Bozoglu, 2016; Zhu et al., 2022). The multifaceted nature of food spoilage has made early identification challenging; however, the ability to evaluate the microbial community as a whole has led to breakthroughs in the identification of spoilage causing organisms and the generation of predictive models to assess risk (Karanth et al., 2023).

As sequencing technologies have improved and become increasingly accessible, 16S rRNA gene sequencing has become a popular tool for researchers aiming to assess the microbial community within an environment (Satam et al., 2023; Weinroth et al., 2022). Despite the widespread application of sequencing-based approaches, it is important to assess limitations relative to the research question. One limitation that poses a major challenge when applying DNA-based technologies to the food industry is the inability to distinguish between DNA from live versus dead cells (Keer & Birch, 2003; Kumar & Ghosh, 2019; Weinroth et al., 2022). Food manufacturers often utilize intervention strategies, such as thermal processing or antimicrobial application, to reduce the microbial population on food products. In these situations, the inability to distinguish between viable and non-viable cellular DNA may result in the misrepresentation of the viable microbial community and the potential false identification of viable pathogenic microorganisms. While there is no perfect solution, a variety of approaches can be utilized to target the viable portion of the microbial population including: RNA-based sequencing, metabolomics, and viability differentiation dyes (i.e. propidium monoazide (PMA) or ethidium monoazide (EMA)) (Chatman et al., 2024; Kumar & Ghosh, 2019; Li et al., 2017). While each technique has its advantages and limitations, viability differentiating dyes have emerged as a cost effective and easy to implement alternative for researchers aiming to remove the non-viable portion of the microbial population prior to molecular quantification (qPCR) or DNA-based sequencing (16S rRNA gene sequencing) (Kumar & Ghosh, 2019; Nocker & Camper, 2006). The efficacy of PMA treatment in combination with PCR to detect and/or quantify different microorganisms has proved promising (Pan & Breidt, 2007; Reyneke et al., 2022). Okada et al. (2022) demonstrated that two-rounds of PMA treatment completely inhibited the detection of non-viable *Campylobacter* spp. in chicken juice by qPCR. Other systematic evaluations have revealed that while PMA does have quantitative potential in simple synthetic communities, its efficacy in complex systems is largely affected by initial biomass, food matrices, and compositional diversity, suggesting further investigation is required (Wang et al., 2021).

The aim of this study was to assess the potential application of 16S rRNA gene sequencing in combination with a viability differentiating dye, PMA, to accurately characterize the microbiota of ready-to-eat (RTE) meat products during manufacturing creating a viable microbial bio-map. The effect of PMA treatment was directly evaluated against a not treated paired control to determine if the removal of non-viable DNA improved sequencing resolution. The utility of the microbial bio-map was assessed based on its ability to identify shifts in the microbial community during manufacturing and the introduction/proliferation of potential spoilage contaminants. Finally, these molecular-based techniques were compared to the more traditional culture-based approach for identifying spoilage organisms in RTE meat products.

## Results

### Sequencing and filtering

A total of 326 samples were sequenced, resulting in 4,968,218 reads and averaging 15,239 reads per sample. After normalization, 38 samples were removed due to low sequencing depth, resulting in 144 Control and 144 PMA samples for further analyses. Only samples with complete pairs were considered when analyzing the treatment effect (PMA vs Control), resulting in a total of 220 samples, 110 pairs.

### Impact of PMA treatment on meat microbiota

Treatment (PMA vs Control) significantly impacted the overall microbial diversity and composition of the paired samples. Samples treated with PMA yielded a less rich microbial community that tended to be less even than the Control group, as determined by α-diversity metrics Observed Features (*p*<0.001; Figure 3a; Supplemental Table 1) and Pielou Evenness (*p*=0.06; Figure 3b; Supplemental Table 1). The impact of treatment was further supported by the observable differences between the two groups (PMA vs Control) in β-diversity metrics, Jaccard Distance (*p*=0.001, *q*=0.001; Figure 3c; Supplemental Table 2) and Weighted UniFrac Distance (*p*=0.001, *q*=0.001; Figure 3d; Supplemental Table 2). A pairwise difference test revealed that PMA did not impact samples equally across all sampling locations (Figure 4). The greatest impact of treatment was observed on raw samples (A) and samples collected in the later stages of processing (i.e. product exiting the holding cooler, sliced product, and packaged product; D-G). More specifically, PMA treatment reduced the richness of the microbial population in raw samples (A; *p*<0.001), samples exiting the holding cooler (D; *p*=0.039), and packaged product samples (G; *p*=0.004). Similarly, PMA treatment reduced the evenness of the microbial community in raw (A; *p*=0.050), sliced (F; *p*=0.003), and packaged product samples (G; *p*=0.003). ANCOM identified three differentially abundant taxonomic groups at the genus level, *Brochothrix* (W=394), *Roseomonas* (W=384), and Enterobacterales order (W=372; Supplemental Figure 1). *Brochothrix* was identified as more abundant in PMA treated samples, while Enterobacterales and *Roseomonas* were detected in a greater abundance in Control samples.

**Figure 1:**
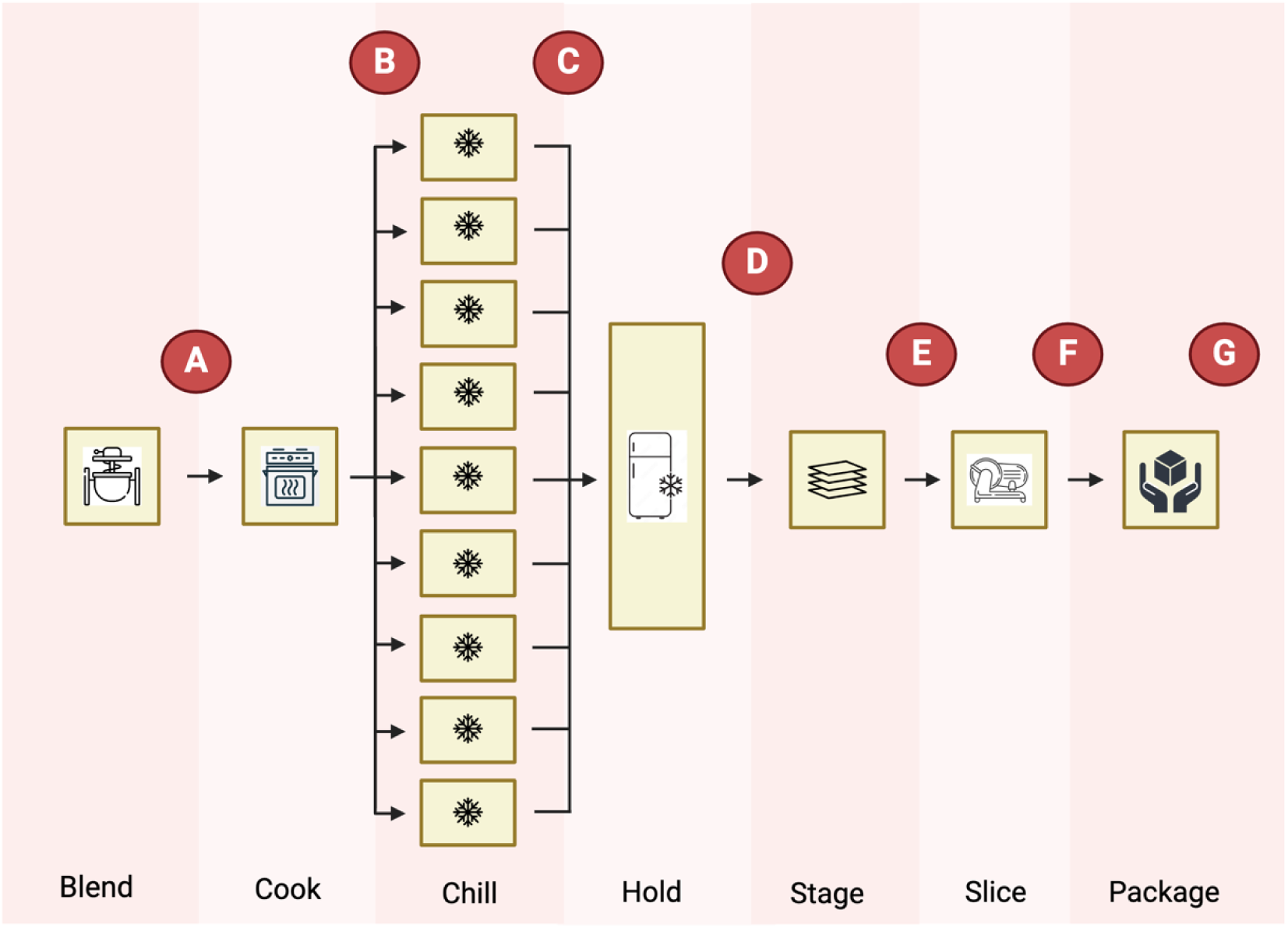
Diagram of the RTE meat product manufacturing flow with sampling locations indicated. Product manufacturing started with the blending of raw materials, followed by thermal processing. The fully cooked product was then chilled in one of nine blast chilling systems and held in a holding cooler, before being staged, sliced, and vacuum packaged. Product sampling locations are indicated by a red circle and labeled according to the following locations: raw material after blending (A), product exiting the oven (B), product exiting the blast chiller (C), product in the holding cooler (D), product in the staging area (E), sliced product (F), and packaged product (G). Nine sets of product samples were collected, following the same batch from start (A) to finish (G), each set representing the use of a blast chiller system. Created in BioRender. Ricke, S. (2025) https://BioRender.com/e49z138.

**Figure 2:**
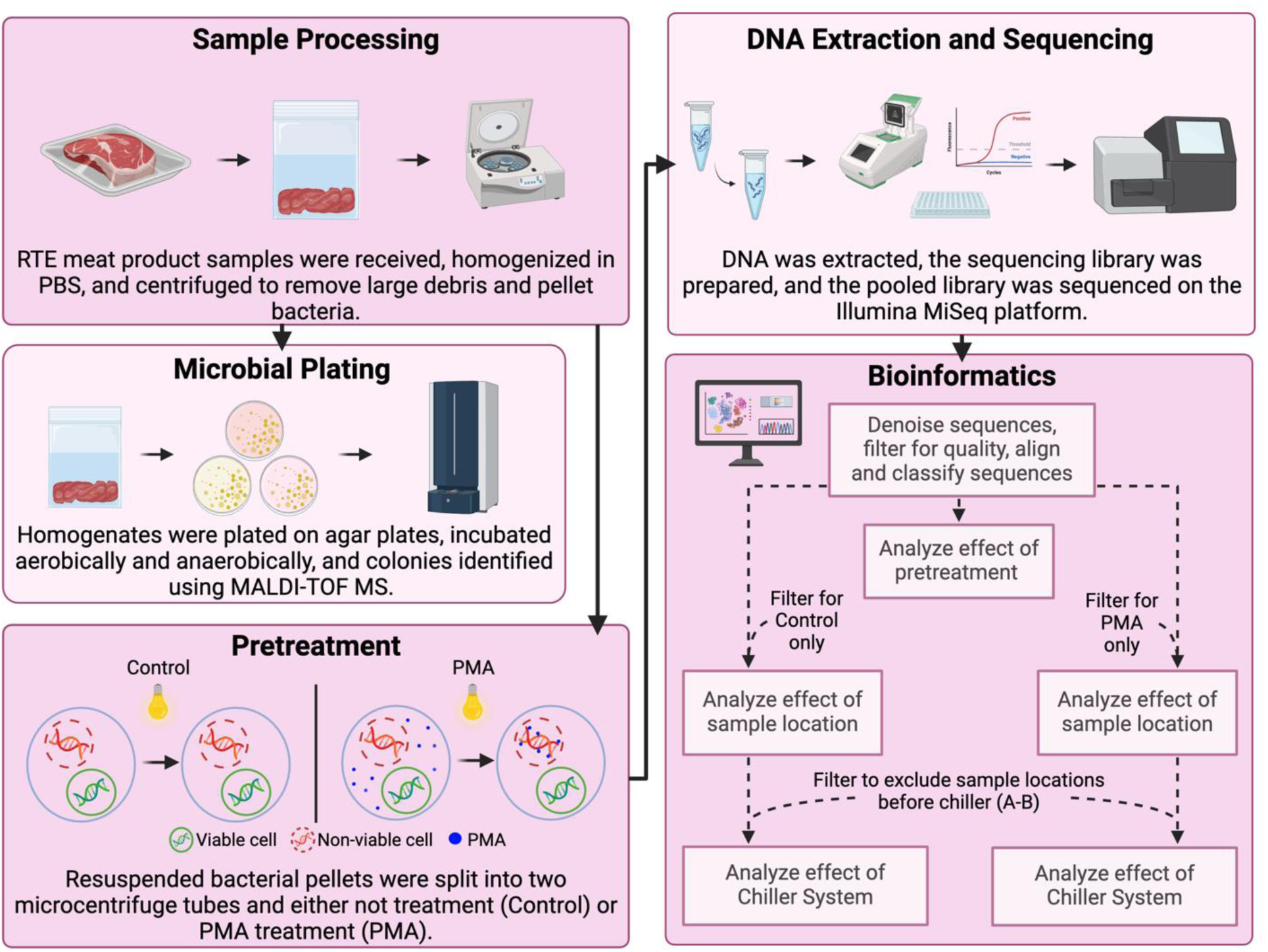
Methodology diagram of sample processing flow from collection to bioinformatics. Samples were subject to all of the same processing steps, with the exception of pretreatment in which samples were subject to either no treatment (Control) or PMA treatment (PMA). Bioinformatic analysis was first performed on the complete dataset to assess pretreatment effect. The dataset was then filtered by pretreatment group (Control vs PMA) and analyzed to determine effect of sampling location. Each of the filtered groups (Control vs PMA) was then filtered again to remove samples collected before exposure to blast chilling systems (A-B) to analyze the effect of chiller system. Created in BioRender. Ricke, S. (2025) https://BioRender.com/9fozgi9.

**Figure 3:**
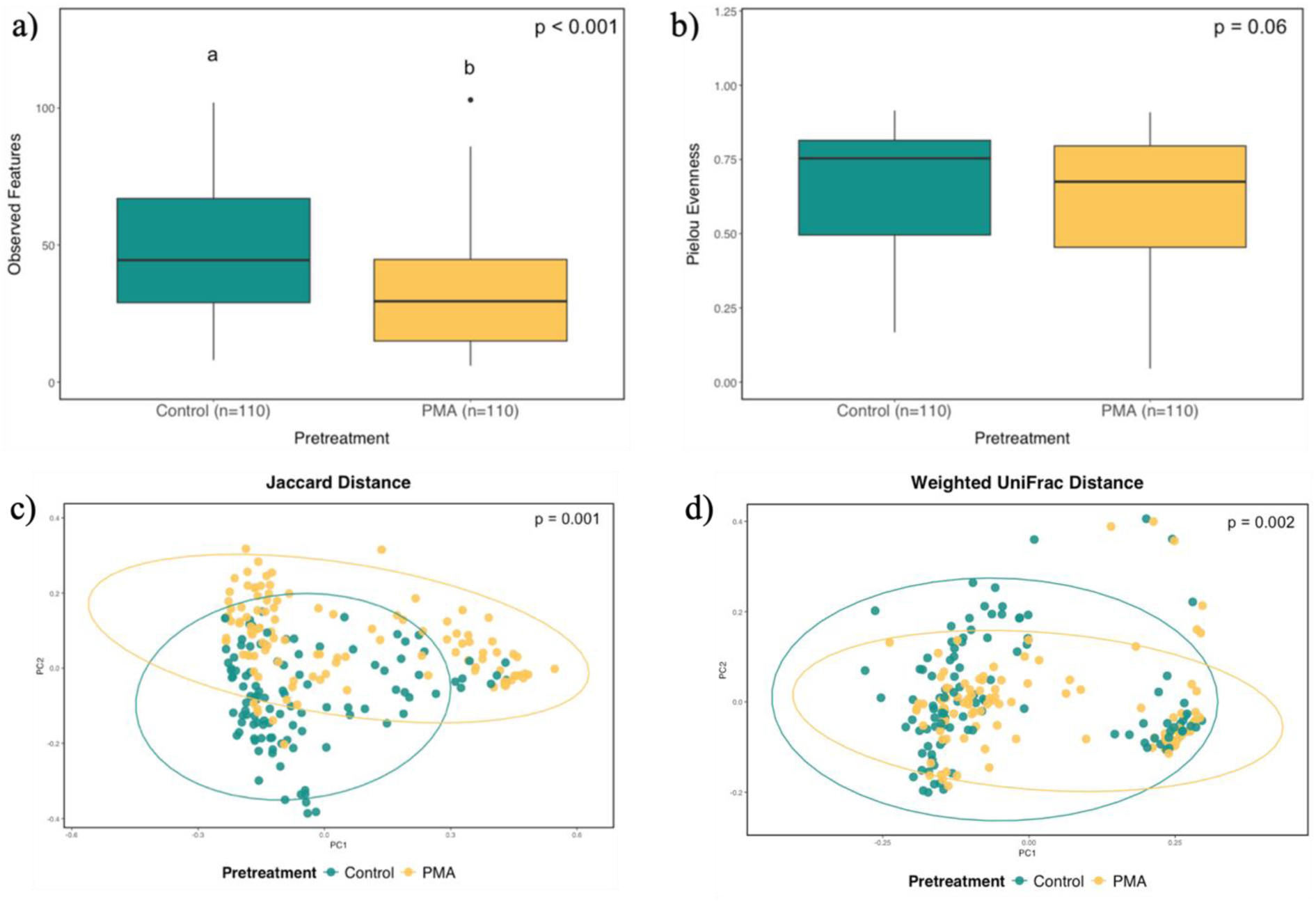
Alpha and beta diversity metrics for RTE meat products by pretreatment. Total genomic DNA was extracted from RTE meat products either not treated (Control) or treated with 25 μM of PMAxx (PMA). Main effect of pretreatment on α-diversity metrics (a) Observed Features and (b) Pielou Evenness were determined using Kruskal-Wallis and pairwise differences were determined using the Wilcoxon signed-rank test. Main effect and pairwise comparisons of β-diversity metrics (c) Jaccard Distance and (d) Weighted UniFrac Distance were determined using ANOSIM. Letters denote pairwise differences between pretreatment (p ≤ 0.05). There was no effect of pretreatment on α-diversity metric Pielou Evenness (*p*=0.06); however, there was an effect of pretreatment on α-diversity metric Observed Features (*p*<0.001) and β-diversity metrics Jaccard Distance (*p*=0.001) and Weighted UniFrac Distance (*p*=0.002) with pairwise differences outlined in Supplemental Table 1 and 2.

**Figure 4:**
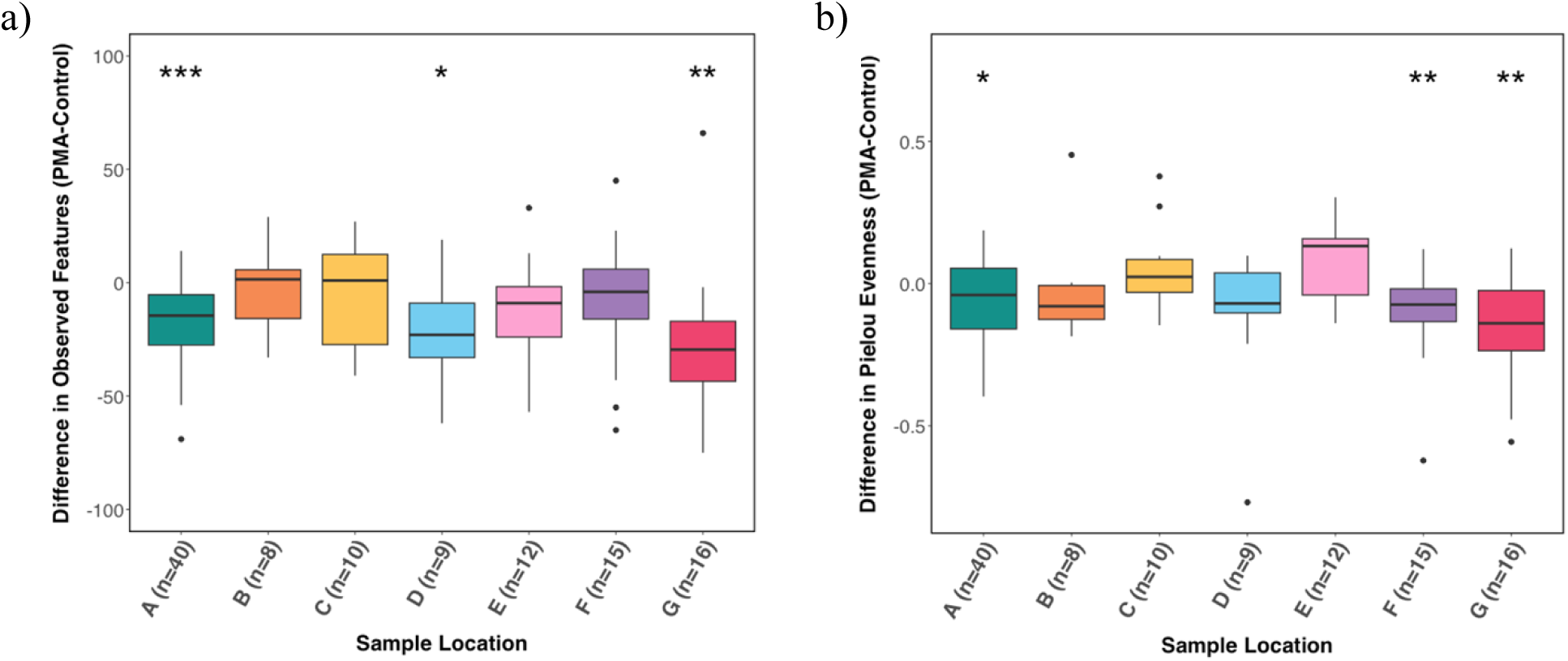
Pairwise differences in alpha diversity metrics for RTE meat products by pretreatment. Total genomic DNA was extracted from RTE meat products either not treated (Control) or treated with 25 μM of PMAxx (PMA). Pairwise difference tests were used to determine the effect of pretreatment, within paired samples, on α-diversity metrics (a) Observed Features and (b) Pielou Evenness. There was an impact of pretreatment on α-diversity metrics Observed Features, for samples collected after raw blending (A), in holding cooler (D), and after packaging (G), and Pielou Evenness, for samples collected after raw blending (A), after slicing (F), and after packaging (G). Asterisks indicate statical significance: * for *p*<0.05, ** for *p*<0.01, *** for *p*<0.001.

### Impact of manufacturing on meat microbiota

In Control samples, one distinct shift in the microbial community was observed after thermal processing (B). As products transitioned from raw material to fully cooked, an increase in microbial richness (Observed Features, *p*<0.001; Figure 5a; Supplemental Table 3) and evenness (Pielou Evenness, *p*<0.001; Figure 5b; Supplemental Table 3) were observed, and subsequently maintained throughout the remainder of manufacturing. Beta-diversity metrics, Jaccard Distance (*p*=0.001; Figure 5c; Supplemental Table 4) and Weighted UniFrac Distance (*p*=0.001; Figure 5d; Supplemental Table 4), also identified that the microbial community of raw samples (A) differed from the community at all other sampling locations (B-G). Thirty-three taxonomic groups were observed to be above the relative abundance threshold of 1% (Figure 6a). The microbial composition of raw product samples (A) consisted primarily of *Pseudomonas*, *Psychrobacter*, *Acinetobacter*, and *Lactobacillus*, making up approximately 46%, 11%, 9%, and 8% of the total population, respectively. These compositional results were further confirmed through the identification of five differentially abundant genera by location: *Acinetobacter* (W=376), *Psychrobacter* (W=375), *Pseudomonas* (W=375), *Myroides* (W=368), *Leuconostoc* (W=356; Supplemental Figure 2). All five genera were determined to be more abundant in raw product samples (A) and nearly absent in samples collected after thermal processing (B-G). *Lactobacillus* was detected throughout manufacturing from raw material (A) to packaged product samples (G), ranging in relative abundance from approximately 8% to 35%; however, it was not identified as being differentially abundant in any specific processing step.

**Figure 5:**
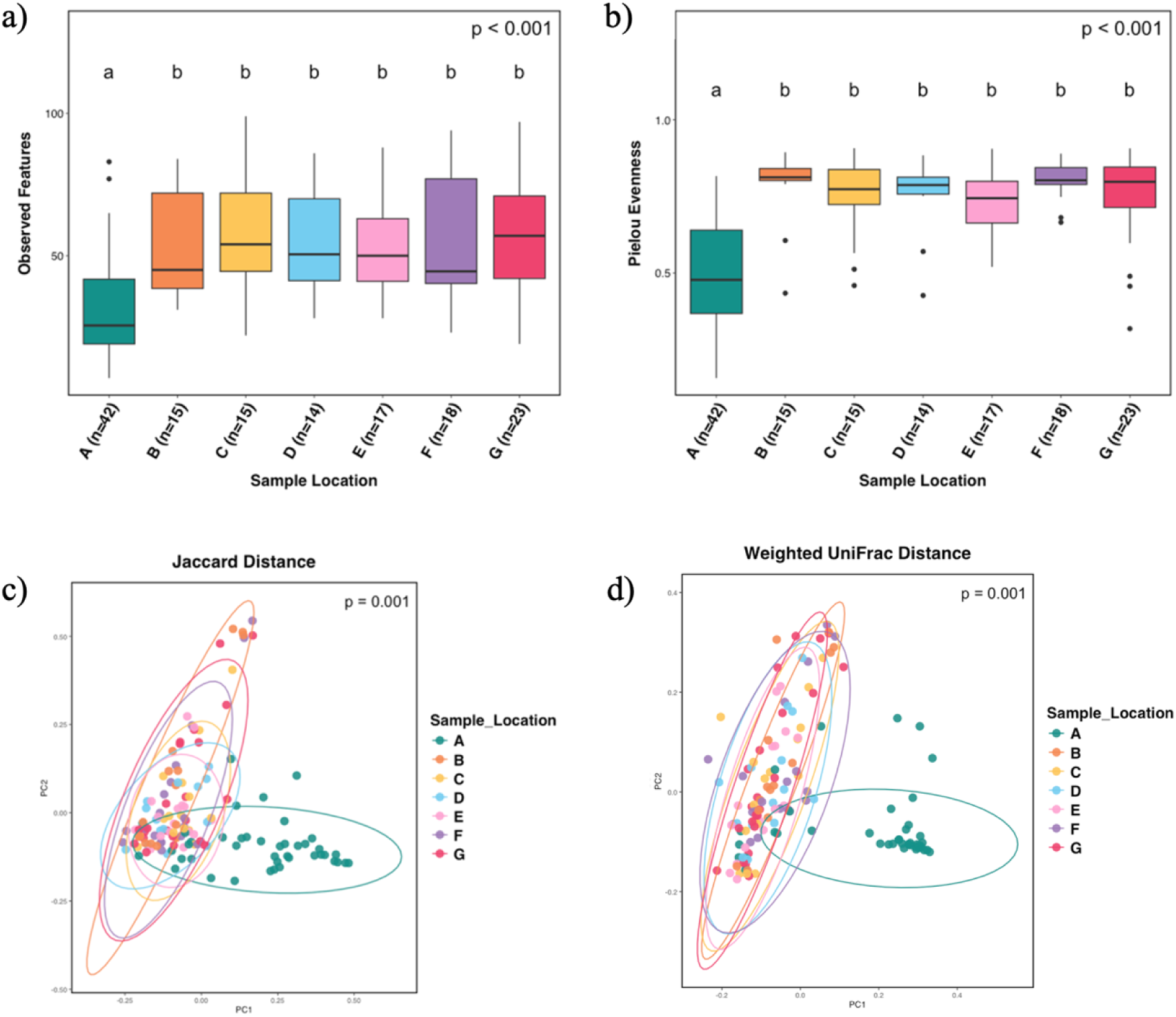
Alpha and beta diversity metrics for not treated (Control) RTE meat products by sample location. Total genomic DNA was extracted from RTE meat products not treated (Control). Main effect of sample location on α-diversity metrics (a) Observed Features and (b) Pielou Evenness were determined using Kruskal-Wallis. Main effect and pairwise comparisons of β-diversity metrics (c) Jaccard Distance and (d) Weighted UniFrac Distance were determined using ANOSIM. Letters denote pairwise differences between treatment (p≤0.05). There was an effect of sample location on α-diversity metrics Observed Features (*p*<0.001) and Pielou Evenness (*p*<0.001), and β-diversity metrics Jaccard Distance (*p*=0.001) and Weighted UniFrac Distance (*p*=0.001) with pairwise differences outlined in Supplemental Table 3 and 4.

**Figure 6:**
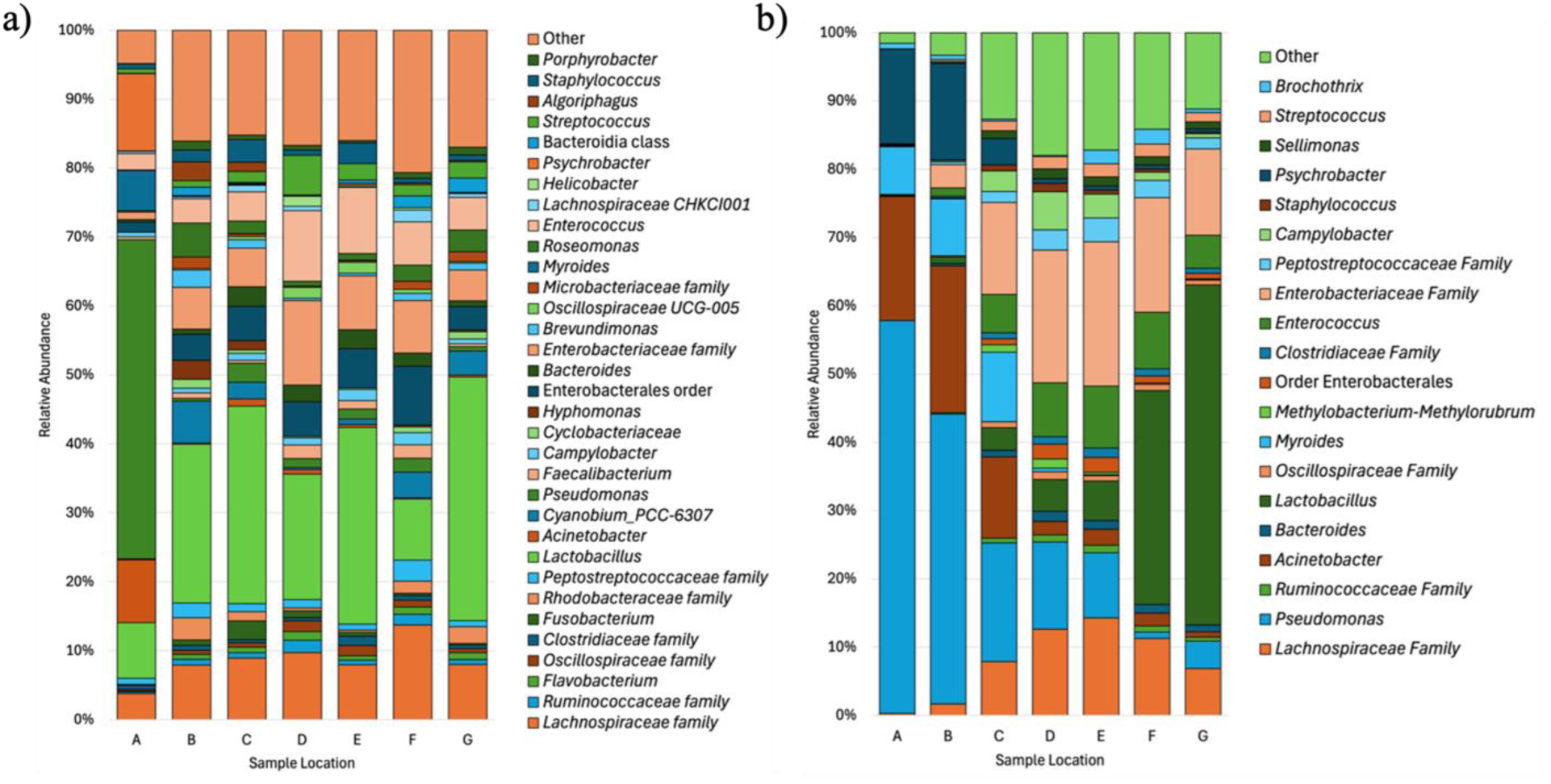
Taxonomic bar plot of the microbial community on not treated (Control) and PMA treated RTE meat products at the genus level. Total genomic DNA was extracted from RTE meat products either not treated (Control; a) or treated with 25 μM of PMAxx (PMA; b). Data was grouped by sample location and represented as mean relative abundances. Bacteria included in figure legend had a relative abundance <1%, all bacteria below this level were grouped into the other category.

In PMA treated samples, two distinct shifts in the microbial community were observed. Similar to the Control group, a shift was identified after thermal processing (B); however, unlike the control group, PMA treatment revealed an additional shift in the microbial population after slicing (F). Alpha-diversity metrics, Observed Features (*p* < 0.001; Figure 7a; Supplemental Table 5) and Pielou Evenness (*p*<0.001; Figure 7b; Supplemental Table 5), identified raw product samples (A) as having the least rich and most uneven microbial population. After the initial increase in microbial richness after thermal processing (B), the number of observed features steadily declined throughout the remainder of manufacturing (B-G). Little variation in community evenness was observed between products exiting the oven (B) and those in the staging area prior to slicing (E); however, after slicing (F) a reduction in evenness was observed. The shift in microbial diversity after slicing (F) was more gradual than the shift observed after thermal processing (B). This gradual reduction in richness can be observed by the lack of significance between samples collected after the holding cooler (D), in the staging area (E), after slicing (F), and packaged product (G; *p*>0.05, *q*>0.05). Beta-diversity metrics, Jaccard Distance (*p* = 0.001; Figure 7c; Supplemental Table 6) and Weighted UniFrac Distance (*p*=0.001; Figure 7d; Supplemental Table 6), displayed a similar trend with slight deviations. Jaccard distance, only taking into consideration community dissimilarities, demonstrated that the raw product samples (A) had the most unique microbial community and had the greatest number of dissimilarities to all other sampling locations (B-G). While some statistical significance was observed between product samples exiting the chiller (C), in the holding cooler (D), after slicing (F), and in the package (G), there remained a large amount of overlap between these locations. When phylogenetic distance and relative abundances were taken into consideration in the Weighted UniFrac Distance metric, three distinct microbial communities were observed, mimicking the shifts observed in α-diversity after thermal processing (B) and slicing (F). These results indicate that while there was overlap in microbial community membership after thermal processing (B), the differences between groups are amplified when relative abundances and phylogenetic distances are taken into consideration. Twenty taxonomic groups at the genus level were identified above the threshold of 1% average relative abundance by sample location (Figure 6b). Similar to the Control group, *Pseudomonas*, *Acinetobacter*, and *Psychrobacter* made up the majority of the microbial composition observed on raw product samples, 57%, 18%, and 14%, respectively (A). Unlike in the Control group, *Lactobacillus* did not emerge as a predominant member of the microbial community until after slicing (F) when the average relative abundance increased from approximately 6% to 31%, and eventually 50% in packaged product (G). Additionally, 11 differentially abundant taxonomic groups were identified at the genus level through ANCOM, including: *Lactobacillus* (W=353), *Pseudomonas* (W=353), *Psychrobacter* (W=353), *Myroides* (W=350), *Lactococcus* (W=349), *Acinetobacter* (W=349), *Enterococcus* (W=344), *Enterobacteriaceae* family (W=353), *Lachnospiraceae* family (W=350), *Ruminococcaceae* family (W=325), and Enterobacterales order (W=340; Supplemental Figure 3). Several of these taxonomic groups, including *Lactococcus*, *Myroides*, *Acinetobacter*, *Pseudomonas*, and *Psychrobacter*, were observed to be more abundant in raw product samples (A). After thermal processing (B), taxonomic groups such as *Enterococcus*, Enterobacterales, *Ruminococcaceae*, *Enterobacteriaceae*, and *Lachnospiraceae*, became more abundant, and finally, the genus *Lactobacillus* was identified as being differentially more abundant in sliced (F) and packaged product (G).

**Figure 7:**
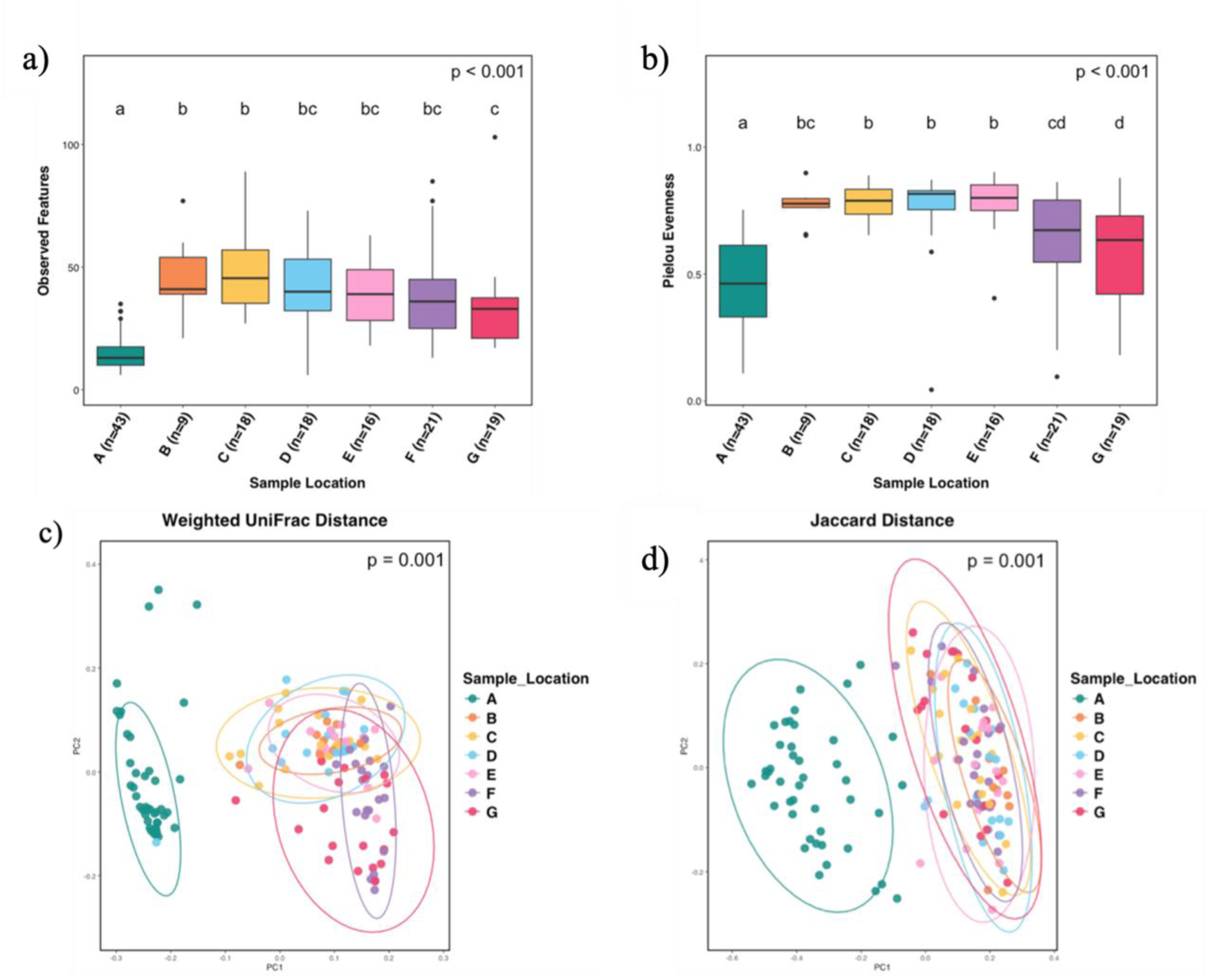
Alpha and beta diversity metrics for PMA treated RTE meat products by sample location. Total genomic DNA was extracted from RTE meat products treated with 25 μM of PMAxx (PMA). Main effect of sample location on α-diversity metrics (a) Observed Features and (b) Pielou Evenness were determined using Kruskal-Wallis. Main effect and pairwise comparisons of β-diversity metrics (c) Jaccard Distance and (d) Weighted UniFrac Distance were determined using ANOSIM. Letters denote pairwise differences between treatment (p≤0.05). There was an effect of sample location on α-diversity metrics Observed Features (*p*<0.001) and Pielou Evenness (*p*<0.001), and β-diversity metrics Jaccard Distance (*p*=0.001) and Weighted UniFrac Distance (*p*=0.001) with pairwise differences outlined in Supplemental Table 5 and 6.

### Impact of chiller system on meat microbiota

Chiller system had a significant impact on the microbial diversity of samples within the Control group. Alpha-diversity metrics, Observed Features (*p* = 0.009; Supplemental Figure 4a; Supplemental Table 7) and Pielou Evenness (p = 0.009; Supplemental Figure 4b; Supplemental Table 7), identified that chiller system had a significant impact on the richness and evenness of the microbial community; however, the only pairwise difference was observed in the richness between samples chilled in chiller systems 3 and 4, with chiller 4 samples possessing a richer microbial community. The effect of chiller system was also observed in β-diversity metrics, Jaccard Distance (p = 0.001; Supplemental Figure 4c; Supplemental Table 8) and Weighted UniFrac Distance (p = 0.001, Supplemental Figure 4d; Supplemental Table 8). Several pairwise differences were observed in the Jaccard Distance metric; however, in the Weighted UniFrac Distance metric only samples chilled in chiller system 8 were identified as being significantly different than those chilled in chiller systems 1, 2, 3, 4, and 9. Three taxonomic groups were identified as differentially abundant by chiller system at the genus level: *Cyanobium* PCC-6307, *Roseomonas*, and the family *Enterobacteriaceae* (Supplemental Figure 5). The genera *Cyanobium* PCC-6307 and *Roseomonas* were identified in a greater abundance on samples chilled in chiller system 7 and 8, while the family *Enterobacteriaceae* was more abundant in samples chilled in chiller system 3.

Individual chiller system had less of an impact on the microbial diversity of PMA treated samples. Richness and evenness of the microbial population were unaffected according to α-diversity metrics, Observed Features (p = 0.379; Supplemental Figure 6a; Supplemental Table 9) and Pielou Evenness (p = 0.673; Supplemental Figure 6b; Supplemental Table 9); however, β-diversity metrics, Jaccard Distance (p = 0.001; Supplemental Figure 6c; Supplemental Table 10) and Weighted UniFrac Distance (p = 0.011; Supplemental Figure 6d; Supplemental Table 10), both exhibited a significant effect of chiller system and, in Jaccard Distance, pairwise differences. Chiller system 3 had the greatest impact on microbial community, as exhibited by the pairwise differences observed between chiller systems 5, 6, 7, and 8. Furthermore, two taxonomic groups at the genus level were identified as being differentially abundant in specific chiller systems. *Acinetobacter* was identified in a greater abundance in samples chilled in chiller systems 1, 2, 3, 4, and 9, while *Pseudomonas* was observed in a greater abundance in samples chilled in chiller systems 4, 5, 7, and 9 (Supplemental Figure 7).

### Culture-based isolation and identification of viable microorganisms

A total of 46 isolates were successfully identified utilizing the MALDI-TOF technique. Twenty-three isolates were collected from raw product samples (A), with the most frequently identified genera being *Bacillus* species (n=6) and *Staphylococcus* (n=3; Figure 8a). Many of these genera were also identified as being present in fully cooked product samples (B-G); however, a number of the isolates collected from cooked product samples were unique to that portion of manufacturing, including *Microbacterium*, *Serratia marcescens*, *Stenotrophomonas*, and *Peribacillus*, to name a few (Figure 8b). *Bacillus* species remained the most frequently isolated microorganism, followed closely by *Enterobacter* species which were isolated at every step of manufacturing (A-G). Finally, two microorganisms were isolated from packaged product (G), Enterobacter and *Lactobacillus sakei*.

**Figure 8:**
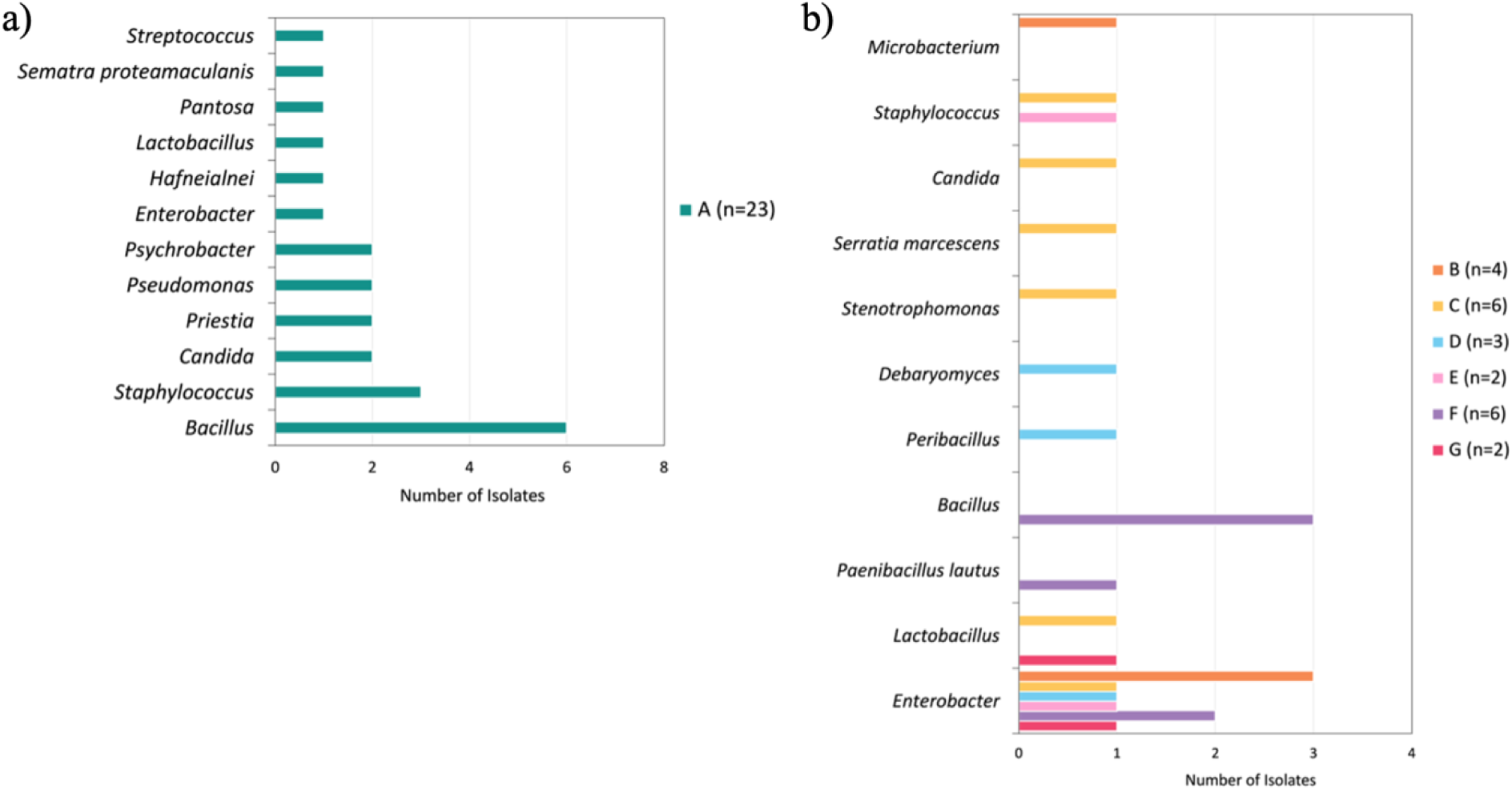
Bacteria cultured from RTE meat product samples and identified using MALDI-TOF MS. Homogenates from raw product (a) and fully cooked product (b) were cultured on TSA, MRS, and APT agar plates under aerobic and anaerobic conditions. Select colonies were isolated based on morphological similarities to known spoilage causing organism and identified using MALDI-TOF MS. Isolates obtaining a score from 2.0 to 2.29 were reported at the genus level, and isolates obtaining a score from 2.3 to 3.0 were reported at the genus and species level. The number of isolates for each location is indicated by the n-value in the figure key, and the number of specific organisms identified is indicated by the scale on the x-axis. *Bacillus* was the most frequently isolated organisms from both raw product and fully cooked product.

## Discussion

In the current study, a viable microbial bio-map was generated by characterizing the microbiota of RTE meat products at various stages of processing in a commercial manufacturing facility. A PMA treatment was applied to a subset of samples at the onset of sample processing to differentiate DNA based on cell membrane integrity and inhibit the amplification of free DNA originating from dead or non-viable cells. The impact of viability differentiation improved the overall resolution of sequencing; however, it was not observed to be equal across all sampling locations, with the greatest impacts being observed at the start and end of manufacturing. The viable portion of the product microbiome experienced two distinct shifts during manufacturing as a result of thermal processing and product slicing, two processing steps that are well documented to impact the microbial community of meat products (Smelt & Brul, 2014; Zagdoun et al., 2020). Our study demonstrated the utility of 16S rRNA gene sequencing in combination with a PMA treatment as a diagnostic tool for RTE meat and poultry processing facilities to identify potential points of contamination and enable the development of targeted, data-driven approaches to improve food safety and quality.

### Diagnostic potential of microbial bio-mapping to understand contamination pathways and environmental influence

The application of high-throughput sequencing in the meat and poultry industry has provided an unparalleled level of resolution into microbial community membership (Imanian et al., 2022). Previous studies have demonstrated the value to be gained from characterizing microbiomes and the versality in which it can be applied (Barcenilla et al., 2024; Bellehumeur et al., 2015; Punchihewage-Don et al., 2025). Poultry production has served as a useful model system to investigate the entire supply chain, demonstrating that the microbial community changes considerably from farm to harvest facility, and again from harvest to processing facility (Park et al., 2023). While longitudinal studies in pork and beef are limited, evidence suggests that these species follow a similar trend in which the final processing environment is a major contributor to final product microbiota (Kang et al., 2019; Y. Kim et al., 2024; Marmion et al., 2021; Park et al., 2023). These studies support our finding that the final RTE meat microbiota is heavily influences by the microbial ecology of the processing environment. Park et al. (2023) demonstrated that proximity plays a key role in the establishment of contamination pathways within a processing facility, as evidenced by the overlapping microbial community membership between food contact surfaces (gloves, worktable, etc.) and final product. Many of the predominant taxonomic groups identified in this study are consistent with those commonly associated with meat processing environments, such as *Pseudomonas*, *Acinetobacter*, and *Psychrobacter* (Barcenilla et al., 2024; Xu et al., 2025). This suggests, that while environmental samples were not collected as a part of this study, there was an interaction between the processing environment and product samples we collected. This relationship was further supported by the observable shift in viable microbial community membership after products came into contact with the slicer which persisted into finished packaged product.

Previous studies have demonstrated that an overall shift in the microbial community occurs during the manufacturing of RTE meat products (Barcenilla et al., 2024; Hultman et al., 2015). While these earlier studies provided insight into the interaction between meat microbiota and processing environment, the lack of intermediate sampling resulted in an inability to evaluate the impact of individual processing steps. In the current study, not only was an overall shift in the microbial community observed from start to finish, similar to existing literature, but through the collection of intermediate samples individual processing steps, like thermal processing and slicing, were identified as having a significant impact on the meat microbiota during product manufacturing.

### The meat microbiota is shaped by individual processing steps and contamination pathways

Thermal processing is a well-documented intervention strategy (Li et al., 2018). In the case of thermally processed RTE meat products, the meat is cooked to an internal temperature of 65 to 71°C to destroy majority of the meat microbiota. It was previously believed that only extremely thermoduric bacteria, such as spore-forming *Bacillus* and *Clostridium* spp., had the ability to survive high heat thermal processing (Erkmen & Bozoglu, 2016). However, more recent evidence suggests that other thermotolerant bacteria such as *Pseudomonas* spp. may also survive traditional thermal processing and grow during chilled storage (Watson et al., 2023). In the current study, the well-documented thermotolerant capabilities of the genus *Lactococcus* may provide some explanation for its persistence from raw to fully cooked product (Kim et al., 2001). The survival of thermotolerant bacteria may account for some of the microbial carryover observed between raw and cooked meat products; however, studies have shown that the final product microbiota is more closely related to the processing environment rather than raw materials, highlighting the contribution of post-lethality contamination pathways (Kim et al., 2024; Xu et al., 2023, 2025).

Meat processing facilities are known to possess persistent bacterial communities, serving as sources of transmission for spoilage and potentially pathogenic microorganisms (Xu et al., 2025). Ready-to-eat products that are sliced and packaged prior to distribution are particularly susceptible to post-lethality contamination and, therefore, must be closely monitored to reduce the risk to public health (Mol et al., 1971). While the risk associated with slicers in retail stores in significantly higher than those in a federally inspected meat processing facility, several studies have confirmed our findings that slicing does impact final product microbiota and potentially reduces the shelf life of presliced meat products (Furbeck et al., 2022; Holley, 1997; USDA FSIS, 2010). Slicers are often regarded as a microbiologically hazardous piece of equipment due to the high potential for surface cross-contamination during use and the difficulty associated with cleaning the equipment (Mertz et al., 2014; Sheen & Hwang, 2010). Our microbial diversity results clearly identified the slicers as a major site of cross-contamination, resulting in a microbial population dominated by *Lactobacillus*. Lactic acid bacteria are commonly found on cooked meat products; therefore, a possible explanation is that *Lactobacillus* from product was transferred to the slicer, it subsequently found a niche and was able to survive sanitation. Similar routes of contamination and persistence have been described for *Pseudomonas* species on slicing equipment in senior living facilities (Koo et al., 2013; Mertz et al., 2014). Furthermore, model systems have indicated the ability of microorganisms to migrate from fermented product to non-fermented product manufactured in the same facility without thorough sanitation in between (Holley, 1997). Being that *Lactobacillus* is commonly used in starter cultures, it is possible that this facility co-manufactures a fermented sausage product resulting in an indigenous population of *Lactobacillus* in the processing environment leading to contamination (Zagorec & Champomier-Vergès, 2017). The isolation of *Lactobacillus sakei* from our finished product samples further supports this potential route of contamination (Figure 8b). Without targeted sampling of the slicer and the processing environment, it is difficult to determine the true source and extent of the contamination; however, the lack of variation observed between collection dates suggests this may be a more persistent contamination. While our study did not focus on the presence of pathogenic microorganisms, these results may serve as an early warning system for insufficient sanitation that could result in a food safety risk.

In addition to the introduction of microorganisms through environmental sources, the microbial composition of the meat microbiota is also heavily influenced by environmental factors such as temperature and atmospheric conditions (Johansson et al., 2020). The predominant taxonomic groups identified in the current study on raw material samples, *Pseudomonas*, *Acinetobacter*, and *Psychrobacter*, were consistent with those commonly observed in other fresh meat studies (Doulgeraki et al., 2012). The chilled storage of fresh meat favors those psychrotrophic microorganisms that are better adapted to survive and thrive at low temperatures, allowing them to outgrow competing members of the microbial community (Ercolini et al., 2009). This phenomenon is likely responsible for the observable low diversity in raw material samples and the steady reduction in microbial richness and evenness observed post-thermal processing. While our study did not follow final product samples after manufacturing, existing literature suggests that this trend in reduced microbial richness will continue throughout the products shelf-life (Dorn-In et al., 2024; Johansson et al., 2020). Modifying atmospheric conditions, either through vacuum packaging or modified atmosphere packaging, is another commonly used method designed to restrict the growth of fast-growing aerobic spoilage microorganisms such as *Pseudomonas* (Borch et al., 1996). The selective pressure enforced by an oxygen depleted environment shifted our predominant bacteria from aerobic Gram-negative microorganisms, observed in the raw material samples, to a microbial community dominated by facultatively anaerobic microorganisms, such as lactic acid bacteria and members of the *Enterobacteriaceae* family. These results are consistent with the existing published literature which recognizes *Lactobacillus*, more specifically *Lactobacillus sakei*, as a dominant member of the microbiota in vacuum packaged chilled pork products and a major contributor to spoilage (Borch et al., 1996; Doulgeraki et al., 2012; Erkmen & Bozoglu, 2016). Spoilage of vacuum packaged RTE meat products by *Lactobacillus* is often characterized by a cloudy appearance and slime formation (Erkmen & Bozoglu, 2016). Although spoiled product was not directly assessed during this study, similar spoilage characteristics were observed prior to the onset of this study, further supporting our theory of *Lactobacillus* induced premature spoilage events.

### Methodological integration improves utility

Two types of methodologies, culture-dependent and -independent, were utilized to investigate the presence of microorganisms during the manufacturing of RTE meat products. This combined approach has been frequently utilized by researchers aiming to capture both quantitative and qualitative information about the microbial community (Dorn-In et al., 2024; Raimondi et al., 2019; Xu et al., 2025). Culture-based techniques have long been used as an efficient and cost-effective way to identify and quantify the viable portion of a microbial community within an environment (Kumar & Ghosh, 2019). Our culture-based approach included the use of selective and non-selective growth mediums to isolate microorganisms present within the product samples and identify them at the genus, and sometimes, species level. This approach provided valuable confirmation of microbial viability and successfully identified potential spoilage microorganisms at various stages of processing. The identification of *Lactobacillus sakei* was the most significant finding, as previously discussed. However, our approach was limited in its ability to accurately represent the overall microbial diversity and provided minimal insight into how the microbial community changed throughout manufacturing.

Culture-independent molecular-based techniques, identify microorganisms based on unique DNA sequences eliminating the need to cultivate viable cells and offers a sensitive, specific, and comprehensive view of the microbial community (Kumar & Ghosh, 2019). Despite these advances, one key limitation of most molecular-based techniques is their inability to distinguish between viable and non-viable cellular DNA(Keer & Birch, 2003; Weinroth et al., 2022; Zeng et al., 2016). This limitation is a challenge for the meat and poultry industry, as extracellular DNA has been shown to survive on surfaces and in solution for extended periods of time, given the right conditions, and the detection of this DNA from non-viable cells can result in the false identification of microorganisms and bias microbial diversity analyses (Arsenault et al., 2024; Dungan et al., 2023; Żarczyńska et al., 2023; Zeng et al., 2016). While a variety of alternative methods are currently used to differentiate viability, none are without limitations (Keer & Birch, 2003; Kumar & Ghosh, 2019). The viability differentiating dye, PMA, provides a promising approach for removing DNA from non-viable cells prior to molecular analysis (Dungan et al., 2023; Miotto et al., 2020; Okada et al., 2022; Reyneke et al., 2022). However, despite initial success, the existing literature recognizes that the efficacy of PMA varies by cellular morphology, microbial complexity, and sample matrix, requiring the need for further, application specific, validation work (Li et al., 2017; Okada et al., 2022; Wang et al., 2021). To our knowledge, our results are the first to demonstrate that the application of a PMA treatment prior to 16S rRNA gene sequencing is an effective strategy to enhance the resolution of microbial analysis in a RTE meat manufacturing setting. Similar results have been obtained when applying the same methodology to dairy products, marine environments, and porcine tissue (Bellehumeur et al., 2015; Mo et al., 2019; Tantikachornkiat et al., 2016).

### Current limitations and taxonomic insights relative to PMA treatment

In this study it was determined that PMA did not impact all sampling locations equally, with the greatest impact being observed on raw material and packaged product samples. Although the proportion of viable and non-viable cells were not quantitatively measured in this study, our results are similar to those previously observed using controlled, live/dead mock communities, supporting the conclusion that the unequal treatment effect was caused by a difference in non-viable cell concentrations (Nocker et al., 2006; Tantikachornkiat et al., 2016). In raw material and packaged product samples, the greater concentration of non-viable DNA is likely due to the indirect consequences of extended storage time and environmental stressors. The removal of the non-viable portion of background DNA revealed an additional shift in the microbial community after slicing that was not distinguishable without PMA treatment. These finding have potential implications for future shelf-life studies, as they suggest sequencing alone may not accurately represent the viable portion of the microbial community of products stored for extended periods of time. Interestingly, PMA exhibited very little impact on the microbial diversity of samples collected immediately following thermal processing, a process known to cause cell death (Smelt & Brul, 2014). High temperatures are known to degrade extracellular DNA, so it is likely that the DNA from non-viable cells was simultaneously released and degraded during thermal process resulting in a low concentration of extracellular DNA (Bauer et al., 2003). Finally, our analysis identified three taxonomic groups as being differentially abundant in either PMA treated (PMA) or not treated (Control) samples: *Brochothrix*, Enterobacterales, and *Roseomonas*. The identification of these taxonomic groups suggests a combination of two hypotheses: (1) PMA is less effective at penetrating Gram-positive cells resulting in a greater abundance of Gram-positive cells, like *Brochothrix*, relative to the remaining portion of the population, and (2) Gram-negative microorganisms, such as Enterobacterales and *Roseomonas*, are more susceptible to processing interventions and/or environmental factors rendering them non-viable and susceptible to PMA treatment. Investigating these theories is beyond the capabilities of this study; however, understanding the mechanisms of PMA and the potential biases it creates will be crucial for future implementation (Wang et al., 2021).

## Conclusion

In conclusion, the results of this study demonstrate the utility of 16S rRNA gene sequencing with viability differentiation as a diagnostic tool for identifying microorganisms during the manufacturing of RTE meat products. To our knowledge this study is the first of its kind to apply a combination of PMA and 16S rRNA gene sequencing to generate a viable microbial bio-map of a commercial RTE meat manufacturing facility. Next-generation sequencing techniques have been previously applied in various segments of the meat industry; however, our results suggest that the use of 16S rRNA gene sequencing alone, may not provide an accurate representation of the viable microbial community, depending on the process interventions being performed. The removal of non-viable DNA through a PMA treatment prior to sequencing proved to be a critical step in improving the resolution of the microbial diversity, especially towards the end of processing. The generation of a comprehensive microbial biomap of a RTE meat manufacturing facility provides the opportunity to identify points of contamination, assess process control, and, with the identification of indicator organisms, may serve as an early detection system for pathogen contamination. The findings from this study will have far reaching implications for the meat and poultry industry relative to experimental design and the interpretation of future results to better account for the presence non-viable cellular DNA.

## Materials and Methods

### Product manufacturing

In this study, product samples were collected from a commercial meat manufacturing facility. The production line of interest produced a cured, ready-to-eat (RTE) comminuted pork product with a final pH of approximately 6.2. The finished product was sliced, vacuum packaged and shipped to retailers fresh. To prevent the growth of pathogenic and spoilage organisms the formulation contained a variety of preservatives, including vinegar, to achieve the desired shelf-life. Manufacturing began with the blending of raw ingredients, followed by thermal processing to an internal temperature of 72°C (162°F), resulting in a fully cooked, RTE meat product. The RTE product was then chilled to 1.6°C (35°F) in one of nine blast chillers and held in a holding cooler at 2.3°C (38°F) for 24 to 96 h. Finally, the product was moved to a staging area at the start of the slicing line where it was sliced and then vacuum packaged (Figure 1).

### Sample collection and processing

Nine sets of product samples were collected at the following seven locations throughout manufacturing: (A) raw blending, (B) after cooking, (C) after chilling, (D) in holding cooler, (E) in staging area, (F) after slicing, (G) after packaging. Each set of samples followed the same batch of product from blending (A) to packaging (G) and represented the use of one blast chiller system. Sample were collected on four individual days within a 2-week period (2/3/23, 2/6/23, 2/13/23, and 2/14/23). At each location, center cut slices were aseptically collected from the top, middle, and bottom of the cart. Samples were stored in sterile bags (Whirl-Pak, Pleasant Prairie, WI, USA) and held at 4°C before being shipped on ice packs to University of Wisconsin-Madison Meat Science and Animal Biologics Discovery (MSABD) laboratory. Upon arrival at the MSABD laboratory, 25 g portions of each sample were transferred to new sterile bags and homogenized by hand with 25 mL of 1× Phosphate Buffered Saline (pH 7.4; PBS; Fisher BioReagents, Waltham, MA, USA). A 10 mL aliquot of the resulting homogenates were transferred to sterile conical tubes (Thermo Scientific, Waltham, MA, USA), while the remainder of the homogenates was used for microbial plating. All conical tubes were subject to a two-step centrifugation process to remove large debris and generate a bacterial pellet. To eliminate large debris, tubes were centrifuged at 800 × g for 10 min at 4°C and supernatants were transferred to new sterile conical tubes. Tubes containing supernatants were then centrifuged at 7000 × g for 10 min to precipitate the bacterial pellet and decanted. The remaining pellets were resuspended in 1 mL of 1× PBS and split between two microcentrifuge tubes (Fisher Scientific, Hampton, USA), containing 500 μL each. These paired samples were then assigned to one of two pretreatment groups, one group was immediately subject to PMA treatment (PMA) while the other group was subject to the same processing steps but remained untreated (Control).

### Microbial plating

Homogenates were spread plated on duplicate sets of Tryptic Soy Agar (TSA; Neogen, Lansing, MI, USA), De Man-Rogosa-Sharpe (MRS; Oxoid, Basingstoke, Hampshire, UK) agar, and All-Purpose Tween (APT; Neogen) agar plates for the cultivation of a wide range of microorganisms including the selective cultivation of homo- and heterofermentative *Lactobacilli*. One set of plates was incubated aerobically at 37 °C for 18 to 24 h and the second set was incubated anaerobically (AnaeroPack^TM^ jar with Anaero Anaerobic Gas Generator sachets; Thermo Scientific) at 37 °C for 18 to 24 to mimic the atmospheric conditions of packaged product. After incubation, plates were examined, and colonies were isolated based on morphological similarities to known spoilage microorganisms. Bacterial colonies were sent to the Wisconsin Veterinary Diagnostic Laboratory (WVDL, Madison, WI, USA) and identified using matrix-assisted laser desorption ionization time of flight mass spectrometry Biotyper 3.1 (MALDI-TOF MS; Bruker Daltonik, Billerica, MA, USA). Bacterial colonies were plated in duplicate onto MALDI-TOF MS polished steel plates. The direct method using formic acid overlayed followed by matrix was performed to extract proteins from the bacteria as described by the manufacturer. Isolates obtaining a score from 2.0 to 2.29 were reported to the genus level, and isolates obtaining a score from 2.3 to 3.0 were reported to the genus and species level.

### PMA treatment

Those samples assigned to the PMA treatment (PMA) group were subject to two-rounds of PMAxx (Biotium, Fremont, CA, USA) treatment prior to DNA extraction. The samples assigned to the not treated (Control) group were also subjected to the same processing steps except 1× PBS was substituted for the PMAxx solution. This treatment was designed to separate the DNA from non-viable cells and prevent amplification in later processing steps. The stock solution of PMAxx was diluted in sterile water to a concentration of 250 μM. Samples were centrifuged at 7000 × g for 10 min, decanted, and resuspended in 180 μL of 1× PBS to concentrate the bacterial pellet. Microcentrifuge tubes containing 180 μL of a bacterial suspension were treated with 20 μL of the 250 μM PMAxx solution to achieve a final concentration of 25 μM PMAxx. Treated samples were vortexed and incubated in the dark at ambient temperatures for 10 min, inverting to mix halfway through. After incubation, samples were placed on a bed of ice and exposed to a 600 W halogen light source (at a distance of 20 cm) for 10 min inverting every 3 to 4 min. After photoactivation, samples were centrifuged at 8,000 × g for 10 min at 4 °C, decanted, and resuspended in 180 μL of 1× PBS. The steps described above were repeated for a total of two-rounds of PMA treatment. After treatment was complete, samples were frozen at -20 °C until DNA extraction could be performed.

### DNA extraction

All samples were subject to DNA extraction using the Qiagen DNeasy Blood and Tissue kit (Qiagen, Hilden, Germany) following the manufacturer protocol developed for tissue samples with slight modifications. Briefly, the frozen samples were thawed, pelleted via centrifugation, and the resulting supernatants discarded. The pellets were resuspended in 180 μL of ATL buffer (Qiagen) and vortexed. Each suspension was subsequently treated with 20 μL of Proteinase K (Qiagen), mixed thoroughly, and incubated in a water bath for 1 h at 56 °C with mixing every 15 min. After incubation, 200 μL of Buffer AL (Qiagen) and 200 μL of molecular grade ethanol (Fisher Scientific, Hampton, USA) were added to each sample and vortexed to mix. The resulting mixtures were then transferred to DNeasy Mini spin columns (Qiagen), positioned in a 2 mL collection tubes, and centrifuged at 6,000 × g for 1 min. The liquid flow through was discarded, and the spin columns were subject to a series of wash steps with AW1 (Qiagen) and AW2 (Qiagen). After the final wash step, the liquid flow through and collection tubes were discarded, and the spin columns were placed in new microcentrifuge tubes. To elute DNA, 50 μL of Buffer AE (Qiagen) was pipetted directly onto the membrane of the spin columns and the columns were incubated at ambient temperatures for 10 min. After incubation, spin columns and microcentrifuge tubes were centrifuged at 6,000 × g for 1 min. After elution, spin columns were discarded and eluted samples were analyzed using a Nanodrop^TM^ One Spectrophotometer (Thermo Fisher Scientific, Waltham, MA, USA) to measure DNA concentrations, 260/280 and 260/230 ratios. Samples with DNA concentrations higher than 15 ng/μL were diluted to 10 ng/μL in Buffer AE (Qiagen) and frozen at -20 °C until library preparation could be performed.

### Library preparation

A sequencing library was prepared using the custom primers designed by Kozich et al. (2013) based on the V4 region of the 16S rRNA gene. Amplification was performed using a high fidelity Accuprime Pfx DNA polymerase (19 μL; Invitrogen, Waltham, MA, USA), dimethyl sulfoxide (2 μL; Fisher Scientific, Hampton, NH, USA), dual-indexed primers containing a unique eight nucleotide barcode sequence (1 μL of 10 mM forward, 1 μL of 10 mM reverse; Integrated DNA Technologies, Coralville, IA, USA), and sample DNA (2 μL). The T100 Thermal Cycler (Bio-Rad Laboratories, Hercules, CA, USA) was set to the following conditions: 95 °C for 5 min, 35 cycles of 95 °C for 30 sec, 55 °C for 30 sec and 72 °C for 1 min, 72 °C for 5 min. Amplification was confirmed using gel electrophoresis (1.5% agarose; Fisher BioReagents). PCR products were normalized to equimolar concentrations using the SequalPrep^TM^ Normalization kit (Invitrogen) and pooled libraries were generated with equal parts of the normalized samples. Pooled library concentrations were determined using a KAPA library quantification kit for Illumina platforms (Roche Sequencing, Indianapolis, IN, USA) and Qubit 1× dsDNA HS Assay Kit (Invitrogen). On the day of sequencing, the pooled library and PhiX Control v3 (Illumina, San Diego, USA) were denatured with freshly made 0.2 N NaOH (Fisher Scientific), incubated for 5 min, and diluted in Hyb buffer (Illumina) to a concentration of 20 pM. The denatured 20 pM pooled library and PhiX were diluted a second time in Hyb buffer (Illumina) to a final concentration of 6 pM. The resulting 6 pM library was combined with the 6 pM PhiX (20%, v/v), loaded into a 2 × 250 cycle v2 MiSeq cartridge (Illumina), and sequenced on an Illumina MiSeq (Illumina). After the sequencing run was complete the resulting sequences were uploaded to BaseSpace (Illumina) and demultiplexed. Demultiplexed reads were uploaded in NCBI (PRJNA1242784) and github (https://github.com/RickeLab-UW/RTE-Meat-Microbiota.git).

### Statistical and bioinformatic analyses

Sequencing data was analyzed using the QIIME2 pipeline (version 2023.9) and RStudio (version 4.4.1; R Core Team, 2024). Demultiplexed paired-end reads were downloaded from Illumina BaseSpace, formatted, and imported into the QIIME2 pipeline using Casava 1.8 (Bolyen et al., 2019). Imported sequences were denoised and filtered for quality with DADA2 via the q2-dada2 plugin (Callahan et al., 2016). Amplicon Sequence Variants (ASVs) were aligned, highly variable positions removed, and phylogenetic trees generated using the align-to-tree-mafft-fasttree pipeline combining commands from the q2-alignment and q2-phylogeny plugins (Katoh & Standley, 2013; Pedregosa et al., 2011; Price et al., 2010; Pruesse et al., 2007; Quast et al., 2012). Taxonomic classification was performed with the q2-feature-classifier plugin using a pretrained SILVA classifier (Silva 138 99% OTUs full-length sequences; UUID: 2bbe61fa-7f78-4913-a6a7-b42e6fff2279) with a specified confidence level of 95% (Bokulich, Kaehler, et al., 2018; Robeson et al., 2021; Rognes et al., 2016). Taxonomy-based filtering was performed using the q2-taxa plugin to exclude features annotated as chloroplast or mitochondria from downstream analysis (McDonald et al., 2012). Metadata-based filtering was also performed to isolate variables such as pretreatment, chiller system, and sampling location (Figure 2). Samples were rarefied to 1,000 reads to remove any residual bias resulting from sequencing depth differences among sample types (n=220, 1,065 to 19,666 frequency/sample) (Caporaso et al., 2011; Villette et al., 2021; Weiss et al., 2017). Diversity analyses were performed through the q2-diversity plugin to generate alpha and beta diversity metrics (McKinney, 2010). The significance of α-diversity metrics, Observed Features (qualitative measure of community richness) and Pielou’s Evenness (measure of community evenness), were determined using Kruskal-Wallis with pairwise differences determined using the Wilcoxon signed-rank test (Kruskal & Wallis, 1952; Pielou, 1966; Weiss et al., 2017). The significance of β-diversity metrics, Jaccard Distance (qualitative measure of community dissimilarity) and Weighted UniFrac Distance (quantitative measure of community dissimilarity that incorporates phylogenetic relationships between features), were determined using analysis of similarities (ANOSIM) (Anderson, 2001; Hamady et al., 2010; Jaccard, 1908; Lozupone et al., 2011; Lozupone et al., 2007; Lozupone & Knight, 2005; McDonald et al., 2018). Pairwise difference tests were used to determine the pretreatment effect of PMA within samples compared to the paired control using the q2-longitudinal plugin (Bokulich, Dillon, et al., 2018). Differential abundance across variables was determined using the analysis of composition of microbiomes (ANCOM) in the q2-composition plugin (Mandal et al., 2015). Main effects and pairwise differences were considered significant at *p* ≤ 0.05 and *q* ≤ 0.05. QIIME2 artifacts were imported into RStudio using the qiime2R package (Bisanz, 2018). Diversity figures and principal coordinates analysis (PCoA) plots were generated in RStudio using ggplot2 package (Wickham, 2016).

## Acknowledgments

The authors would like to thank our industry collaborator for providing commercial samples and financial support throughout this study. Author J.A.B. would like to acknowledge the University of Wisconsin Foundation at the University of Wisconsin-Madison for its continued support through the Bray-Woodbury Wisconsin Distinguished Graduate Fellowship Fund in Meat Science. Authors are grateful to Ashley Tarcin, Margaret Costello, Grace Drexel, and Rachel Ganske for their valuable assistance during data collection. The authors declare that the research was conducted in the absence of any commercial or financial relationship that could be construed as a potential conflict of interest.

## Literature Cited

Anderson, MJ. 2001. A new method for non-parametric multivariate analysis of variance. Austral Ecology, 26(1), 32–46. 10.1111/j.1442-9993.2001.01070.pp.x

Arsenault H, Kuffel A, Daeid NN, Gray A. 2024. Trace DNA and its persistence on various surfaces: A long term study investigating the influence of surface type and environmental conditions – Part one, metals. Forensic Science International: Genetics, 70, 103011. 10.1016/j.fsigen.2024.103011

Barcenilla C, Cobo-Díaz JF, Puente A, Valentino V, De Filippis F, Ercolini D, Carlino N, Pinto F, Segata N, Prieto M, López M, Alvarez-Ordóñez A. 2024. In-depth characterization of food and environmental microbiomes across different meat processing plants. Microbiome, 12(1), 199. 10.1186/s40168-024-01856-3

Bauer T, Weller P, Hammes WP, Hertel C. 2003. The effect of processing parameters on DNA degradation in food. European Food Research and Technology, 217(4), 338–343. 10.1007/s00217-003-0743-y

Bellehumeur C, Boyle B, Charette SJ, Harel J, L’Homme Y, Masson L, Gagnon CA. 2015. Propidium monoazide (PMA) and ethidium bromide monoazide (EMA) improve DNA array and high-throughput sequencing of porcine reproductive and respiratory syndrome virus identification. Journal of Virological Methods, 222, 182–191. 10.1016/j.jviromet.2015.06.014

Bisanz JE. 2018. qiime2R: Importing QIIME2 artifacts and associated data into R sessions (v0.99). https://github.com/jbisanz/qiime2R

Bokulich NA, Dillon MR, Zhang Y, Rideout JR, Bolyen E, Li H, Albert PS, Caporaso JG. 2018. q2-longitudinal: Longitudinal and paired-sample analyses of microbiome data. MSystems, 3(6). 10.1128/msystems.00219-18

Bokulich NA, Kaehler BD, Rideout JR, Dillon M, Bolyen E, Knight R, Huttley GA, Caporaso JG. 2018. Optimizing taxonomic classification of marker-gene amplicon sequences with QIIME 2’s q2-feature-classifier plugin. Microbiome, 6(1), 90. 10.1186/s40168-018-0470-z

Bolyen E, Rideout JR, Dillon MR, Bokulich NA, Abnet CC, Al-Ghalith GA, Alexander H, Alm EJ, Arumugam M, Asnicar F, Bai Y, Bisanz JE, Bittinger K, Brejnrod A, Brislawn CJ, Brown CT, Callahan BJ, Caraballo-Rodríguez AM, Chase J, … Caporaso JG. 2019. Reproducible, interactive, scalable and extensible microbiome data science using QIIME 2. Nature Biotechnology, 37(8), 852–857. 10.1038/s41587-019-0209-9

Borch E, Kant-Muermans MLT, Blixt Y. 1996. Bacterial spoilage of meat and cured meat products. International Journal of Food Microbiology, 33(1), 103–120. 10.1016/0168-1605(96)01135-X

Buzby JC, Wells HF, Hyman J. 2014. The estimated amount, value, and calories of postharvest food losses at the retail and consumer levels in the United States. www.ers.usda.gov/publications/eib-economic-information-bulletin/EIB-121.aspx

Callahan BJ, McMurdie PJ, Rosen MJ, Han AW, Johnson AJA, Holmes SP. 2016. DADA2: High-resolution sample inference from Illumina amplicon data. Nature Methods, 13(7), 581–583. 10.1038/nmeth.3869

Caporaso JG, Lauber CL, Walters WA, Berg-Lyons D, Lozupone CA, Turnbaugh PJ, Fierer N, Knight R. 2011. Global patterns of 16S rRNA diversity at a depth of millions of sequences per sample. Proceedings of the National Academy of Sciences, 108(supplement_1), 4516–4522. 10.1073/pnas.1000080107

Chatman CC, Olson EG, Mantovani HC, Majumder EL, Ricke SC. 2024. Meat animal biologics discovery opportunities from the gut microbiome: application of metabolomics. Meat and Muscle Biology, 8(1). 10.22175/mmb.18261

Dorn-In S, Mang S, Cosentino RO, Schwaiger K. 2024. Changes in the microbiota from fresh to spoiled meat, determined by culture and 16S rRNA analysis. Journal of Food Protection, 87(2), 100212. 10.1016/j.jfp.2023.100212

Doulgeraki AI, Ercolini D, Villani F, Nychas G-J. 2012. Spoilage microbiota associated to the storage of raw meat in different conditions. International Journal of Food Microbiology, 157(2), 130–141. 10.1016/j.ijfoodmicro.2012.05.020

Dungan AM, Geissler L, Williams AS, Gotze CR, Flynn EC, Blackall LL, van Oppen MJH. 2023. DNA from non-viable bacteria biases diversity estimates in the corals *Acropora loripes* and *Pocillopora acuta*. Environmental Microbiome, 18(1), 86. 10.1186/s40793-023-00541-6

Ercolini D, Russo F, Nasi A, Ferranti P, Villani F. 2009. Mesophilic and psychrotrophic bacteria from meat and their spoilage potential *in vitro* and in beef. Applied and Environmental Microbiology, 75(7), 1990–2001. 10.1128/AEM.02762-08

Erkmen O, Bozoglu TF. 2016. Spoilage of meat and meat products. In Food Microbiology: Principles into Practice (1st ed., pp. 279–295). Wiley. 10.1002/9781119237860.ch16

Furbeck RA, Bower CG, Fernando SC, Sullivan GA. 2022. A survey of the microbial communities of commercial presliced, packaged deli-style ham throughout storage. Meat and Muscle Biology, 6(1). 10.22175/mmb.15446

Grunert KG. 2005. Food quality and safety: consumer perception and demand. European Review of Agricultural Economics, 32(3), 369–391. 10.1093/eurrag/jbi011

Hamady M, Lozupone CA, Knight R. 2010. Fast UniFrac: facilitating high-throughput phylogenetic analyses of microbial communities including analysis of pyrosequencing and PhyloChip data. The ISME Journal, 4(1), 17–27. 10.1038/ismej.2009.97

Holley RA. 1997. Impact of slicing hygiene upon shelf life and distribution of spoilage bacteria in vacuum packaged cured meats. Food Microbiology, 14(3), 201–211. 10.1006/fmic.1996.0089

Hultman J, Rahkila R, Ali J, Rousu J, Björkroth KJ. 2015. Meat processing plant microbiome and contamination patterns of cold-tolerant bacteria causing food safety and spoilage risks in the manufacture of vacuum-packaged cooked sausages. Applied and Environmental Microbiology, 81(20), 7088–7097. 10.1128/AEM.02228-15

Imanian B, Donaghy J, Jackson T, Gummalla S, Ganesan B, Baker RC, Henderson M, Butler EK, Hong Y, Ring B, Thorp C, Khaksar R, Samadpour M, Lawless KA, MacLaren-Lee I, Carleton HA, Tian R, Zhang W, Wan J. 2022. The power, potential, benefits, and challenges of implementing high-throughput sequencing in food safety systems. npj Science of Food, 6(1), 35. 10.1038/s41538-022-00150-6

Jaccard P. 1908. Nouvelles Recherches Sur La Distribution Floral. Bull. Soc. Vard. Sci. Nat, 44, 223–270.

Johansson P, Jääskeläinen E, Nieminen T, Hultman J, Auvinen P, Björkroth KJ. 2020. Microbiomes in the context of refrigerated raw meat spoilage. Meat and Muscle Biology, 4(2). 10.22175/mmb.10369

Kang S, Ravensdale J, Coorey R, Dykes GA, Barlow R. 2019. A Comparison of 16S rRNA profiles through slaughter in Australian export beef abattoirs. Frontiers in Microbiology, 10. 10.3389/fmicb.2019.02747

Karanth S, Feng S, Patra D, Pradhan AK. 2023. Linking microbial contamination to food spoilage and food waste: the role of smart packaging, spoilage risk assessments, and date labeling. Frontiers in Microbiology, 14. 10.3389/fmicb.2023.1198124

Katoh K, Standley DM. 2013. MAFFT multiple sequence alignment software version 7: improvements in performance and usability. Molecular Biology and Evolution, 30(4), 772–780. 10.1093/molbev/mst010

Keer JT, Birch L. 2003. Molecular methods for the assessment of bacterial viability. Journal of Microbiological Methods, 53(2), 175–183. 10.1016/S0167-7012(03)00025-3

Kim WS, Ren J, Dunn NW. 2001. Assessment of the tolerance of *Lactococcus lactis* cells at elevated temperatures. Biotechnology Letters, 23, 1141–1145.

Kim Y, Ban G-H, Hong YW, Jeong KC, Bae D, Kim SA. 2024. Bacterial profile of pork from production to retail based on high-throughput sequencing. Food Research International, 176, 113745. 10.1016/j.foodres.2023.113745

Koo OK, Mertz AW, Akins EL, Sirsat SA, Neal JA, Morawicki R, Crandall PG, Ricke SC. 2013. Analysis of microbial diversity on deli slicers using polymerase chain reaction and denaturing gradient gel electrophoresis technologies. Letters in Applied Microbiology, 56(2), 111–119. 10.1111/lam.12021

Kozich JJ, Westcott SL, Baxter NT, Highlander SK, Schloss PD. 2013. Development of a dual-index sequencing strategy and curation pipeline for analyzing amplicon sequence data on the MiSeq Illumina sequencing platform. Applied and Environmental Microbiology, 79(17), 5112–5120. 10.1128/AEM.01043-13

Kruskal WH, Wallis WA. 1952. Use of ranks in one-criterion variance analysis. Journal of the American Statistical Association, 47(260), 583–621. 10.1080/01621459.1952.10483441

Kumar SS, Ghosh AR. 2019. Assessment of bacterial viability: a comprehensive review on recent advances and challenges. Microbiology, 165(6), 593–610. 10.1099/mic.0.000786

Li R, Kou X, Zhang L, Wang S. 2018. Inactivation kinetics of food-borne pathogens subjected to thermal treatments: a review. International Journal of Hyperthermia, 34(2), 177–188. 10.1080/02656736.2017.1372643

Li R, Tun HM, Jahan M, Zhang Z, Kumar A, Dilantha Fernando WG, Farenhorst A, Khafipour E. 2017. Comparison of DNA-, PMA-, and RNA-based 16S rRNA Illumina sequencing for detection of live bacteria in water. Scientific Reports, 7(1), 5752. 10.1038/s41598-017-02516-3

Lozupone CA, Hamady M, Kelley ST, Knight R. 2007. Quantitative and qualitative β diversity measures lead to different insights into factors that structure microbial communities. Applied and Environmental Microbiology, 73(5), 1576–1585. 10.1128/AEM.01996-06

Lozupone CA, Knight R. 2005. UniFrac: a new phylogenetic method for comparing microbial communities. Applied and Environmental Microbiology, 71(12), 8228–8235. 10.1128/AEM.71.12.8228-8235.2005

Lozupone CA, Lladser ME, Knights D, Stombaugh J, Knight R. 2011. UniFrac: an effective distance metric for microbial community comparison. The ISME Journal, 5(2), 169–172. 10.1038/ismej.2010.133

Mandal S, Van Treuren W, White RA, Eggesbø M, Knight R, Peddada SD. 2015. Analysis of composition of microbiomes: a novel method for studying microbial composition. Microbial Ecology in Health & Disease, 26(0). 10.3402/mehd.v26.27663

Marmion M, Ferone MT, Whyte P, Scannell AGM. 2021. The changing microbiome of poultry meat; from farm to fridge. Food Microbiology, 99, 103823. 10.1016/j.fm.2021.103823

McDonald D, Clemente JC, Kuczynski J, Rideout JR, Stombaugh J, Wendel D, Wilke A, Huse S, Hufnagle J, Meyer F, Knight R, Caporaso JG. 2012. The biological observation matrix (BIOM) format or: how I learned to stop worrying and love the ome-ome. GigaScience, 1(1), 7. 10.1186/2047-217X-1-7

McDonald D, Vázquez-Baeza Y, Koslicki D, McClelland J, Reeve N, Xu Z, Gonzalez A, Knight R. 2018. Striped UniFrac: enabling microbiome analysis at unprecedented scale. Nature Methods, 15(11), 847–848. 10.1038/s41592-018-0187-8

McKinney, W. (2010). Data structures for statistical computing in Python. In S. van der Walt & J. Millman (Eds.), 9th Python in Science Conference (pp. 51–56).

Mertz AW, Koo OK, O’Bryan CA, Morawicki R, Sirsat SA, Neal JA, Crandall PG, Ricke SC. 2014. Microbial ecology of meat slicers as determined by denaturing gradient gel electrophoresis. Food Control, 42, 242–247. 10.1016/j.foodcont.2014.02.027

Miotto M, Barretta C, Ossai SO, da Silva HS, Kist A, Vieira CRW, Parveen S. 2020. Optimization of a propidium monoazide-qPCR method for *Escherichia coli* quantification in raw seafood. International Journal of Food Microbiology, 318, 108467. 10.1016/j.ijfoodmicro.2019.108467

Mo L, Yu J, Jin H, Hou Q, Yao C, Ren D, An X, Tsogtgerel T, Zhang H. 2019. Investigating the bacterial microbiota of traditional fermented dairy products using propidium monoazide with single-molecule real-time sequencing. Journal of Dairy Science, 102(5), 3912–3923. 10.3168/jds.2018-15756

Mol JHH, Hietbrink JEA, Mollen HWM, Van Tinteren J. 1971. Observations on the microflora of vacuum packed sliced cooked meat products. Journal of Applied Bacteriology, 34(2), 377–397. 10.1111/j.1365-2672.1971.tb02297.x

Nocker A, Camper AK. 2006. Selective removal of DNA from dead cells of mixed bacterial communities by use of ethidium monoazide. Applied and Environmental Microbiology, 72(3), 1997–2004. 10.1128/AEM.72.3.1997-2004.2006

Nocker A, Cheung C-Y, Camper AK. 2006. Comparison of propidium monoazide with ethidium monoazide for differentiation of live vs. dead bacteria by selective removal of DNA from dead cells. Journal of Microbiological Methods, 67(2), 310–320. 10.1016/j.mimet.2006.04.015

Okada A, Tsuchida M, Rahman MM, Inoshima Y. 2022. Two-round treatment with propidium monoazide completely inhibits the detection of dead *Campylobacter* spp. cells by quantitative PCR. Frontiers in Microbiology, 13. 10.3389/fmicb.2022.801961

Pan Y, Breidt F. 2007. Enumeration of viable *Listeria monocytogenes* cells by real-time PCR with propidium monoazide and ethidium monoazide in the presence of dead cells. Applied and Environmental Microbiology, 73(24), 8028–8031. 10.1128/AEM.01198-07

Park J, Bae D, Kim SA. 2023. Microbial trace investigation throughout the entire chicken supply chain based on metagenomic high-throughput sequencing. Food Research International, 169, 112775. 10.1016/j.foodres.2023.112775

Pedregosa F, Varoquaux G, Gramfort A, Michel V, Thirion B, Grisel O, Blondel M, Prettenhofer P, Weiss R, Dubourg V, Vanderplas J, Passos A, Cournapeau D, Brucher M, Perrot M, Duchesnay É. 2011. Scikit-learn: Machine learning in Python. Journal of Machine Learning Research, 12, 2825–2830. http://scikit-learn.sourceforge.net.

Petrescu DC, Vermeir I, Petrescu-Mag RM. 2019. Consumer understanding of food quality, healthiness, and environmental impact: A cross-national perspective. International Journal of Environmental Research and Public Health, 17(1), 169. 10.3390/ijerph17010169

Pielou EC. 1966. The measurement of diversity in different types of biological collections. Journal of Theoretical Biology, 13, 131–144. 10.1016/0022-5193(66)90013-0

Price MN, Dehal PS, Arkin AP. 2010. FastTree 2 – Approximately maximum-likelihood trees for large alignments. PLoS ONE, 5(3), e9490. 10.1371/journal.pone.0009490

Pruesse E, Quast C, Knittel K, Fuchs BM, Ludwig W, Peplies J, Glockner FO. 2007. SILVA: a comprehensive online resource for quality checked and aligned ribosomal RNA sequence data compatible with ARB. Nucleic Acids Research, 35(21), 7188–7196. 10.1093/nar/gkm864

Punchihewage-Don AJ, Hasan NA, Parveen S. 2025. 16S rRNA amplicon sequencing of organic and conventional chickens. Microbiology Resource Announcements. 10.1128/mra.01081-24

Quast C, Pruesse E, Yilmaz P, Gerken J, Schweer T, Yarza P, Peplies J, Glöckner FO. 2012. The SILVA ribosomal RNA gene database project: improved data processing and web-based tools. Nucleic Acids Research, 41(D1), D590–D596. 10.1093/nar/gks1219

R Core Team. 2024. R: A language and environment for statistical computing. R Foundation for Statistical Computing. https://www.R-project.org/

Raimondi S, Luciani R, Sirangelo TM, Amaretti A, Leonardi A, Ulrici A, Foca G, D’Auria G, Moya A, Zuliani V, Seibert TM, Søltoft-Jensen J, Rossi M. 2019. Microbiota of sliced cooked ham packaged in modified atmosphere throughout the shelf life. International Journal of Food Microbiology, 289, 200–208. 10.1016/j.ijfoodmicro.2018.09.017

Reyneke B, Waso M, Ndlovu T, Clements T, Havenga B, Khan S, Khan W. 2022. EMA-versus PMA-amplicon-based sequencing to elucidate the viable bacterial community in rainwater. Water, Air, & Soil Pollution, 233(4), 103. 10.1007/s11270-022-05578-w

Robeson MS, O’Rourke DR, Kaehler BD, Ziemski M, Dillon MR, Foster JT, Bokulich NA. 2021. RESCRIPt: Reproducible sequence taxonomy reference database management. PLOS Computational Biology, 17(11), e1009581. 10.1371/journal.pcbi.1009581

Rognes T, Flouri T, Nichols B, Quince C, Mahé F. 2016. VSEARCH: a versatile open source tool for metagenomics. PeerJ, 4(10), e2584. 10.7717/peerj.2584

Satam H, Joshi K, Mangrolia U, Waghoo S, Zaidi G, Rawool S, Thakare RP, Banday S, Mishra AK, Das G, Malonia SK. 2023. Next-generation sequencing technology: Current trends and advancements. Biology, 12(7), 997. 10.3390/biology12070997

Sheen S, Hwang C-A. 2010. Mathematical modeling the cross-contamination of *Escherichia coli* O157:H7 on the surface of ready-to-eat meat product while slicing. Food Microbiology, 27(1), 37–43. 10.1016/j.fm.2009.07.016

Smelt JPPM, Brul S. 2014. Thermal inactivation of microorganisms. Critical Reviews in Food Science and Nutrition, 54(10), 1371–1385. 10.1080/10408398.2011.637645

Tantikachornkiat M, Sakakibara S, Neuner M, Durall DM. 2016. The use of propidium monoazide in conjunction with qPCR and Illumina sequencing to identify and quantify live yeasts and bacteria. International Journal of Food Microbiology, 234, 53–59. 10.1016/j.ijfoodmicro.2016.06.031

USDA FSIS. 2010. FSIS Comparative risk assessment for Listeria monocytogenes in ready-to-eat meat and poultry deli meats report.

van Herpen E, de Hooge IE. 2019. When product attitudes go to waste: Wasting products with remaining utility decreases consumers’ product attitudes. Journal of Cleaner Production, 210, 410–418. 10.1016/j.jclepro.2018.10.331

Villette R, Autaa G, Hind S, Holm JB, Moreno-Sabater A, Larsen M. 2021. Refinement of 16S rRNA gene analysis for low biomass biospecimens. Scientific Reports, 11(1), 10741. 10.1038/s41598-021-90226-2

Wang Y, Yan Y, Thompson KN, Bae S, Accorsi EK, Zhang Y, Shen J, Vlamakis H, Hartmann EM, Huttenhower C. 2021. Whole microbial community viability is not quantitatively reflected by propidium monoazide sequencing approach. Microbiome, 9(1), 17. 10.1186/s40168-020-00961-3

Watson SC, Furbeck RA, Fernando SC, Chaves BD, Sullivan GA. 2023. Spoilage *Pseudomonas* survive common thermal processing schedules and grow in emulsified meat during extended vacuum storage. Journal of Food Science, 88(5), 2162–2167. 10.1111/1750-3841.16557

Weinroth MD, Belk AD, Dean C, Noyes N, Dittoe DK, Rothrock MJ, Ricke SC, Myer PR, Henniger MT, Ramírez GA, Oakley BB, Summers KL, Miles AM, Ault-Seay TB, Yu Z, Metcalf JL, Wells JE. 2022. Considerations and best practices in animal science 16S ribosomal RNA gene sequencing microbiome studies. Journal of Animal Science, 100(2). 10.1093/jas/skab346

Weiss S, Xu ZZ, Peddada S, Amir A, Bittinger K, Gonzalez A, Lozupone CA, Zaneveld JR, Vázquez-Baeza Y, Birmingham A, Hyde ER, Knight R. 2017. Normalization and microbial differential abundance strategies depend upon data characteristics. Microbiome, 5(1), 27. 10.1186/s40168-017-0237-y

Wickham H. 2016. ggplot2: Elegant graphics for data analysis. Springer-Verlag New York. https://ggplot2.tidyverse.org

Xu ZS, Ju T, Yang X, Gänzle M. 2023. A meta-analysis of bacterial communities in food processing facilities: Driving forces for assembly of core and accessory microbiomes across different food commodities. Microorganisms, 11(6), 1575. 10.3390/microorganisms11061575

Xu ZS, Pham VD, Yang X, Gänzle MG. 2025. High-throughput analysis of microbiomes in a meat processing facility: are food processing facilities an establishment niche for persisting bacterial communities? Microbiome, 13(1), 25. 10.1186/s40168-024-02026-1

Zagdoun M, Coeuret G, N’Dione M, Champomier-Vergès M-C, Chaillou S. 2020. Large microbiota survey reveals how the microbial ecology of cooked ham is shaped by different processing steps. Food Microbiology, 91, 103547. 10.1016/j.fm.2020.103547

Zagorec M, Champomier-Vergès M-C. 2017. *Lactobacillus sakei*: A starter for sausage fermentation, a protective culture for meat products. Microorganisms, 5(3), 56. 10.3390/microorganisms5030056

Żarczyńska M, Żarczyński P, Tomsiao M. 2023. Nucleic acids persistence—Benefits and limitations in forensic genetics. Genes, 14(8), 1643. 10.3390/genes14081643

Zeng D, Chen Z, Jiang Y, Xue F, Li B. 2016. Advances and challenges in viability detection of foodborne pathogens. Frontiers in Microbiology, 7(NOV). 10.3389/fmicb.2016.01833

Zhu Y, Wang W, Li M, Zhang J, Ji L, Zhao Z, Zhang R, Cai D, Chen L. 2022. Microbial diversity of meat products under spoilage and its controlling approaches. Frontiers in Nutrition, 9. 10.3389/fnut.2022.1078201

